# Perceptual sensitivity supports online control, while perceptual prediction errors drive motor memory formation during locomotor adaptation

**DOI:** 10.64898/2026.08.01.742204

**Authors:** Marcela Gonzalez-Rubio, Amber R. Costello, Pablo A. Iturralde, Gelsy Torres-Oviedo

## Abstract

To maintain stable locomotion, the nervous system must continually adapt, whether reacting to an unexpected perturbation, such as a trip on uneven terrain, or anticipating external demands, such as walking on snow, by forming and updating motor memories. Error-based learning is the dominant computational account of such adaptation, in which motor commands are updated to reduce prediction errors, that is, the mismatch between predicted and actual limb state. Yet this framework has been largely defined in reduced, single-effector paradigms, where the sensory consequences of movement are isolated and the prediction error is directly controlled by the experimenter. Whether the same principle governs whole-body, multi-segmental, multi-sensory behaviors such as walking has remained untested, owing in part to the challenge of identifying a behavioral proxy for prediction errors in this complex, dynamic setting. Here, we combined a split-belt treadmill paradigm with a novel method for quantifying perception of leg motion to address this open question. We found two perceptual contributions to locomotor adaptation: perceptual sensitivity predicted initial motor corrections during early adaptation, whereas perceptual prediction errors (i.e., the mismatch between the actual belt-speed difference and the difference perceived as symmetric) predicted the magnitude of motor aftereffects during post-adaptation. Together, these findings support error-based learning, long characterized in upper limb tasks, as a core computational principle underlying human locomotion.

**Significance:** Error-based learning is the dominant account of how the brain adapts movement, yet it has been defined almost entirely in reduced laboratory tasks, such as cursor rotations, planar reaching, and saccades, where prediction errors are directly controlled by the experimenter. Whether it governs whole-body behaviors such as walking has remained untested. Using a novel measure of leg-motion perception during split-belt walking, we show that two dimensions of perception contribute separably to adaptation. Perceptual sensitivity, a proxy for sensory uncertainty, relates to real-time motor corrections, consistent with Bayesian integration. Perceptual prediction error, a proxy for the prediction error that underlies error-based learning, relates to motor memory formation. These findings establish leg-motion perception as a behavioral window onto the computations driving locomotor adaptation, providing a framework for understanding individual and clinical variability in motor behavior.

## 3 Introduction

Whether reacting to an unexpected perturbation, such as a trip on uneven terrain, or anticipating specific external demands, such as walking on snow, by forming and updating motor memories, the nervous system must continually adapt to maintain stable locomotion. When it encounters a novel environment, two complementary processes restore movement (Morton and Bastian, 2006; Shadmehr et al., 2010; Seethapathi et al., 2024). A feedback controller uses incoming sensory signals to detect deviations from expected movement and generate reactive corrections in real time; these occur quickly, require no practice, and are not stored. A memory-based feedforward controller generates anticipatory motor commands from prior experience, requiring practice and producing a stored movement pattern that persists as aftereffects once the perturbation is removed (Morton and Bastian, 2006; Seethapathi et al., 2024). A central question is what computational principle drives adaptation in both.

The dominant answer is error-based learning driven by prediction errors. As the brain sends motor commands, it predicts their sensory consequences via internal models, which are representations that map motor commands onto expected states of the body and environment. The mismatch between these predictions and incoming sensory information constitutes a prediction error, thought to drive real-time feedback corrections and, when sustained, to update the internal models behind feedforward commands, giving rise to motor memories (Wolpert et al., 1995; Shadmehr et al., 2010). This framework, however, was defined almost entirely in reduced, single-effector paradigms, such as visuomotor cursor rotations, planar reaching, and saccadic eye movements, which made the theory tractable by isolating the sensory consequences of movement, so that the perturbation imposed by the experimenter could directly control the prediction error. Whether the same principle governs whole-body, multi-sensory, dynamic behaviors such as walking has remained largely untested.

Establishing whether error-based learning governs walking is challenging because prediction errors are difficult to isolate in a multi-segmental, multi-sensory, dynamic behavior, where, unlike in reduced tasks, the imposed perturbation cannot be assumed to directly control them. Moreover, error-based learning may not be the operative principle: performance-based accounts propose instead that locomotor adaptation optimizes movement outcomes such as energy consumption and stability, with sensory signals informing a reinforcement process rather than directly encoding prediction errors (Seethapathi et al., 2024; Finley et al., 2013; Sánchez et al., 2019). These accounts are not mutually exclusive, but they assign different roles to sensory signals. Under error-based learning, prediction errors directly drive recalibration, whereas under performance-based optimization, sensory signals serve to evaluate movement outcomes. Testing the error-based account therefore requires a behavioral measure of the prediction errors themselves.

The discrepancy between the actual state of the body and the individual’s internal reference of it, which is behaviorally accessible through perceptual tasks, may represent the prediction error driving learning. We will call this discrepancy “perceptual prediction error”. In reaching, perceptual prediction errors reflect the updating of internal models (Zhang et al., 2024, 2026), such that individuals with larger perceptual prediction errors show greater motor aftereffects (Tsay et al., 2022; Zhang et al., 2026). Split-belt walking shifts both perceived step lengths (Sombric et al., 2019) and perceived leg-speed symmetry (Vazquez et al., 2015; Statton et al., 2018; Leech et al., 2018; Rossi et al., 2024, 2026), confirming that shifts in this internal reference occur in this context. Critically, both perception (Vazquez et al., 2015; Leech et al., 2018) and motor output (Iturralde and Torres-Oviedo, 2019) show aftereffects that reverse the direction of the original exposure once the perturbation is removed. This parallel recalibration raises the question of whether the two are functionally linked, yet the evidence remains contested, with some studies reporting independence (Statton et al., 2018; Leech et al., 2018) and others suggesting coupling (Rossi et al., 2026). Whether error-based learning genuinely governs locomotor adaptation has thus been assumed more than demonstrated. We therefore used perception of leg motion as a behavioral window onto the prediction error signal, testing whether perceptual prediction errors in walking relate both to motor corrections during adaptation and to motor aftereffects during post-adaptation.

Perception, however, is not fully characterized by perceptual prediction errors alone. Perceptual sensitivity, the precision with which sensory differences can be discriminated, constitutes a second, functionally distinct dimension that reflects the level of sensory uncertainty (Ernst and Banks, 2002) and thus serves as a proxy for the reliability weighting central to Bayesian integration. Two questions follow. First, whether perceptual sensitivity adapts alongside perceived leg-speed symmetry or remains unchanged: in reaching, sensitivity consistently does not change following visuomotor or force-field adaptation, even as perceived hand position shifts substantially (Cressman and Henriques, 2009; Salomonczyk et al., 2013; Kitchen and Miall, 2021; Ostry et al., 2010), but whether this dissociation extends to walking is unknown. Second, whether sensitivity plays a functional role. Bayesian integration proposes that greater sensory uncertainty should attenuate the influence of sensory-mediated errors, such as prediction errors, on recalibration, because uncertain signals are down-weighted relative to prior expectations (Körding and Wolpert, 2004). Empirical evidence is scarce and mixed: reaching studies find no relationship between sensitivity and motor changes during or after adaptation (Kitchen and Miall, 2021; Vandevoorde and Orban de Xivry, 2021), whereas a whole-body study found that better sensitivity to the direction of body motion improves balance during challenging beam walking (Mirdamadi et al., 2024), suggesting sensitivity may aid real-time corrections. Whether perceptual sensitivity predicts motor changes during adaptation or post-adaptation in walking remains to be tested.

Here we addressed these questions using split-belt walking as a model of sensorimotor adaptation in a whole-body, multi-sensory, dynamic task, combining it with a novel method for quantifying perception of leg motion throughout adaptation (Gonzalez-Rubio et al., 2025, 2026). We assessed two dimensions of perception and related each to motor behavior. First, motivated by error-based learning, we characterized how perception adapts and tested whether individuals with larger perceptual prediction errors show greater motor aftereffects. Second, motivated by Bayesian integration, we examined whether perceptual sensitivity adapts and tested whether greater sensitivity led to more efficient corrections and stronger motor memories. Together, these tests ask whether error-based learning, long characterized in reduced tasks, generalizes to human locomotion.

## 4 Results

### Perceptual prediction errors and motor corrections are reduced during split-belt adaptation

To investigate how perception and motor outputs adapt during split-belt walking, fourteen neurotypical young adults (N=14; 9 females, 20.86 *±* 1.92 y.o.) experienced a period of asymmetric (split-belt) walking in which one belt moved faster than the other (500 mm/s), preceded by a baseline tied-belt period and followed by a post-adaptation tied-belt period. Throughout the experiment, participants performed a two-alternative forced choice (2AFC) task (perceptual probes, Fig 1A), in which they judged which leg felt slower (Gonzalez-Rubio et al., 2025, 2026), allowing us to track how their perception of leg-speed differences evolved alongside their motor patterns.

**Figure 1:**
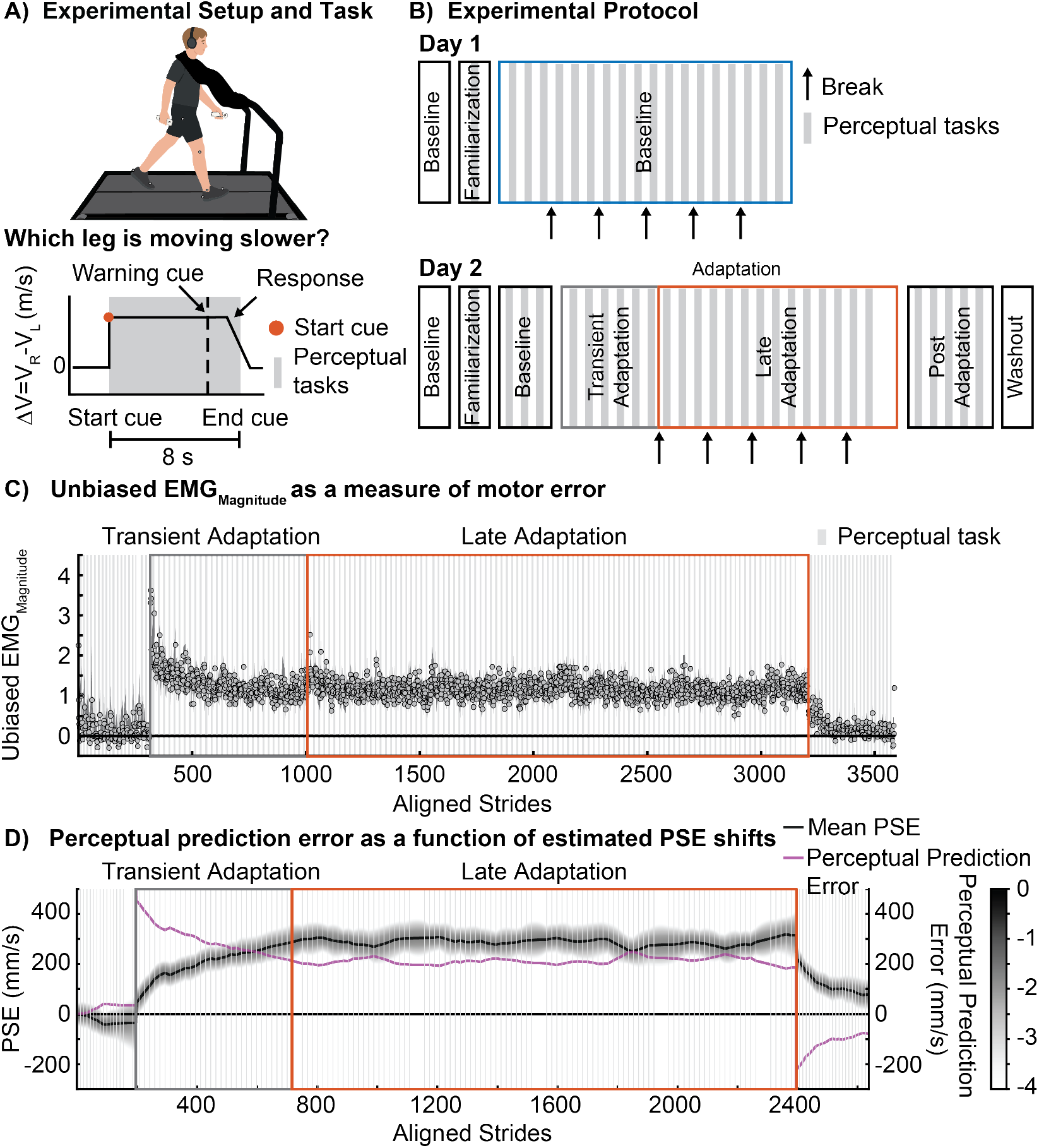
Experimental design and group-level timecourses. **A)** Experimental setup (top) and perceptual task schematic (bottom). Participants walked on a split-belt treadmill while performing a two-alternative forced choice (2AFC) task in which they judged which leg felt slower. A speed difference (Δ*V*) was introduced as a stimulus during an 8 second decision window, marked by an auditory cue. Participants responded by pressing a button in the handheld controller to indicate which leg felt slower, and received auditory confirmation of their response. No feedback on response accuracy was provided. **B)** Experimental protocol across two sessions. Session 1 consisted of three blocks of tied-belt walking with interleaved perceptual probes to establish baseline perception. Session 2 included a brief baseline block followed by the Adaptation period, divided into Transient and Late phases, during which perceptual probes were interleaved with sustained split-belt walking, and a Post-adaptation period in which perceptual probes were interleaved with tied-belt walking. **C)** EMG_Magnitude_ timecourse. Each point represents the average normalized summed muscle activity across 10 bilateral lower-leg muscles at each gait cycle, expressed relative to baseline. Gray and orange rectangle indicate Transient and Late adaptation. Gray shaded areas indicate the perceptual tasks. **D)** Perceptual predictions error and group-level PSE timecourse. PSE probability density (gray) and group mean PSE (black) are shown alongside perceptual prediction error (purple). The PSE trajectory was estimated through a Hidden Markov Model, tracking the continuous evolution of perceived symmetry across strides.

Motor changes were quantified as the overall magnitude of lower-limb muscle activity (EMG_Magnitude_; see Methods), and used as a measure of motor output. As expected, EMG_Magnitude_ rose sharply at the onset of split-belt walking, reflecting the immediate neuromuscular demand of the perturbation, and decreased progressively through Late Adaptation as participants restored efficient motor patterns (Fig 1C). Upon returning to tied-belt walking, EMG_Magnitude_ initially resembled the activity level from the perturbed environment before returning to baseline, consistent with established motor aftereffect dynamics.

Perception of limb speed adapted over the same protocol, following a previously uncharacterized time course. To reconstruct the continuous trajectory of perceived limb speed, we fitted a Hidden Markov Model (HMM) to group-level responses on the 2AFC task across the adaptation protocol, yielding a stride-by-stride estimate of the point of subjective equality (PSE; the speed difference perceived as symmetric, in mm/s) over the full experiment (Fig 1D). This provided the dynamics of perceptual adaptation at a resolution not previously available in walking studies, which have typically assessed PSE at discrete preand post-adaptation time points. The PSE can be read as the participant’s internal reference for “normal”, the belt-speed difference that feels symmetric at any given moment, and its distance from the actual belt-speed difference (0 mm/s during baseline and post-adaptation, 500 mm/s during adaptation) defines the perceptual prediction error, capturing how far the current experience departs from what feels normal (Fig 1D).

During baseline, both belts moved at the same speed and the PSE sat near zero, so the symmetric environment felt normal and the perceptual prediction error was small. At the onset of the 500 mm/s split perturbation, the PSE remained near its baseline value: participants’ sense of normal had not yet changed, so the imposed limb-speed asymmetry was perceived as atypical, yielding a large perceptual prediction error. As adaptation progressed, the PSE shifted steadily toward the imposed difference: the internal reference migrated toward the sustained perturbation, which increasingly felt normal, so that the perceptual prediction error decreased, plateauing by late adaptation. During post-adaptation, the belts returned to equal speeds but the adapted PSE persisted. In other words, the newly established sense of normal limb-speed persisted, so that a zero speed difference now felt atypical, a perceptual aftereffect.

Because both perceptual error and motor output were reduced during adaptation and showed aftereffects, we asked whether participants with larger residual perceptual errors showed larger motor aftereffects.

### Perceptual prediction errors are associated with motor aftereffects, but not motor corrections during split-belt adaptation

While the Hidden Markov model captured the continuous group-level trajectory of perceptual adaptation, we used a Generalized Linear Mixed-Effects Model (GLMM) to obtain individual-level estimates at discrete epochs, enabling us to quantify how much each participant’s perceptual reference shifted and to test its relationship with motor outcomes. We described the probability of choosing “left is slower” as a function of the speed difference for each walking epoch (Fig 2A): 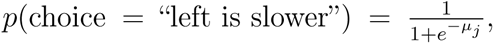 with *µ_j_* = *β*_0_*_j_* + *β*_1_*_j_*Δ*V* + *β*_2_*_j_Epoch* + *β*_3_*_j_*Δ*V* : *Epoch*, where *j* is each participant, *Epoch* is either Baseline (blue) or Late Adaptation (orange), and *β*_2_ and *β*_3_ represents the difference in intercept and slope between the epochs, respectively. From these functions, we estimated the PSE, and computed perceptual prediction error as the difference between the actual belt speed difference (500 mm/s) and changes in PSE (Fig 2B). This quantity captures the systematic discrepancy between what participants perceive as symmetric (bias) and what the belts are actually doing. A large perceptual prediction error indicates that the internal symmetry reference has shifted little toward the imposed perturbation, whereas a small error reflects substantial perceptual recalibration.

**Figure 2:**
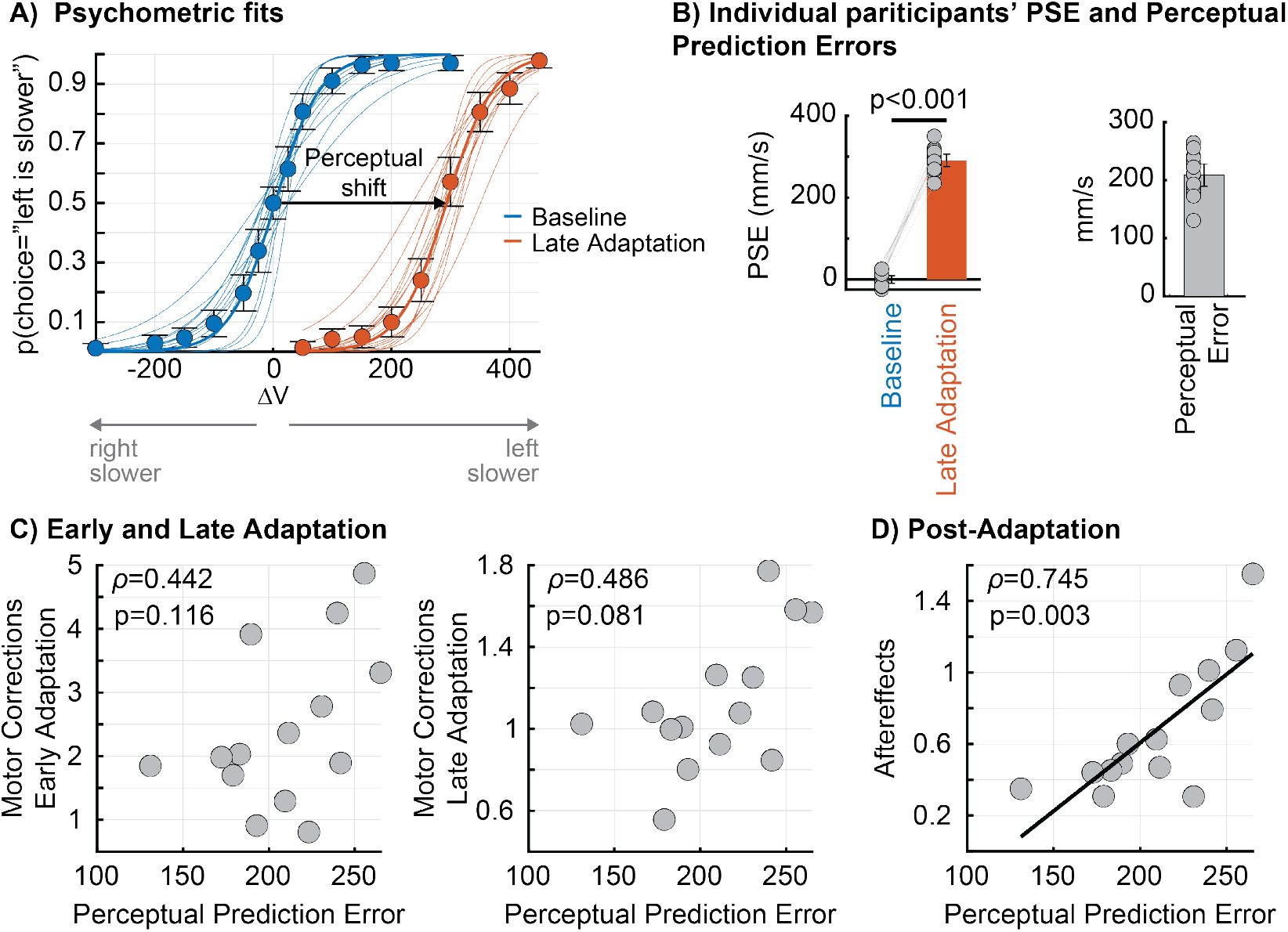
Perceptual adaptation and its relationship to motor outcomes. **A)** Psychometric functions fitted to individual participants’ 2AFC responses during Baseline (blue) and Late Adaptation (orange). Thin lines represent individual participants’ fitted psychometric functions and thick lines represent the group average. Circles represent pooled proportions of ‘left is slower’ responses across participants and error bars represent *±* 1.96 SE. The rightward shift of the group psychometric function by Late Adaptation indicates that participants changed their perception of symmetric walking after adaptation. **B)** Individual participants’ PSE (left) and residual perceptual prediction error (right) at Baseline (blue) and Late Adaptation (orange). Bar height represents the group mean, gray circles represent individual participants, lines connect each participant’s values across epochs, error bar represents *±* 1.96 SE, and horizontal black lines indicate significant differences between epochs assessed by paired t-test. **C)** Spearman correlations between residual perceptual prediction error at Late Adaptation and motor corrections during Early Adaptation (left) and Late Adaptation (right). Gray circles represent individual participants. **D)** Spearman correlation between residual perceptual prediction error at Late Adaptation and motor aftereffects during Post-adaptation. Gray circles represent individual participants. Regression line is shown to indicate significant correlation.

Split-belt walking produced a robust rightward shift in the psychometric function at late adaptation (Fig 2A, orange curve), confirmed by a significant change in the model intercept (*β̄*_2_ = *−*7.321*, p <* 0.001). The PSE shifted from 0.173 *±* 17.447 mm/s at Baseline to 291.222 *±* 28.805 mm/s during Late Adaptation (Fig 2B, left panel; *p <* 0.001). Individual PSE shifts ranged from 235 mm/s to 369 mm/s (47-74% of the actual speed difference experienced of 500 mm/s).

We next examined whether residual perceptual prediction errors, as a proxy for prediction errors, were associated with motor corrections during adaptation and motor aftereffects during post-adaptation, as predicted by error-based learning, which posits that larger prediction errors drive stronger updating of internal models and consequently greater motor corrections and aftereffects. Contrary to this prediction, we only found a robust positive association between perceptual prediction error and motor aftereffects (Fig 2D, right panel; *ρ* = 0.745*, p* = 0.003), but perceptual prediction error was not significantly associated with motor corrections during early or late adaptation ( Early Adaptation, Fig 2C, left panel; *ρ* = 0.442*, p* = 0.116; Late Adaptation (Fig 2C, right panel; *ρ* = 0.486*, p* = 0.081). In other words, participants who experienced larger perceptual prediction errors exhibited larger motor aftereffects upon return to tied-belt walking. These results suggest that perceptual prediction errors are selectively linked to motor memory formation rather than real-time motor corrections. This pattern was not observed when using step length asymmetry as the motor outcome (perceptual prediction errors vs. step length asymmetry during Early Adaptation *ρ* = 0.402*, p* = 0.155; Late Adaptation *ρ* = 0.459*, p* = 0.101; or Post-adaptation *ρ* = *−*0.244*, p* = 0.4), consistent with previous reports (Iturralde and Torres-Oviedo, 2019).

### Perceptual sensitivity remains stable during split-belt adaptation

Beyond changes in perceptual prediction errors, participants may also change their perceptual sensitivity, which refers to the precision with which interlimb leg-speed differences are detected. This is quantified as the Just Noticeable Difference (JND) and serves as a behavioral index of sensory uncertainty. The JND can be visualized with the slope of the psychometric function fitted to perceptual responses. We re-plotted the psychometric functions during Baseline (blue) and Late Adaptation (orange) aligned to their respective PSE values, enabling a direct visual comparison of slopes across epochs (Fig 3A). The slopes appeared to be similar between Baseline and Late Adaptation, suggesting that the precision of speed discrimination was unaffected by splitbelt walking. This visual observation was confirmed statistically. The model revealed no significant interaction between stimulus magnitude and epoch (*β̄*_3_ = *−*0.001*, p* = 0.625), indicating that the slope of the psychometric functions, and therefore perceptual sensitivity, did not change from Baseline to Late Adaptation. Correspondingly, JND values did not differ significantly between epochs (paired t-test, Fig 3B; *p* = 0.236). Thus, while prolonged exposure to split-belt walking induced robust changes in perceptual prediction errors, it did not alter the precision with which participants could detect speed differences between their legs.

**Figure 3:**
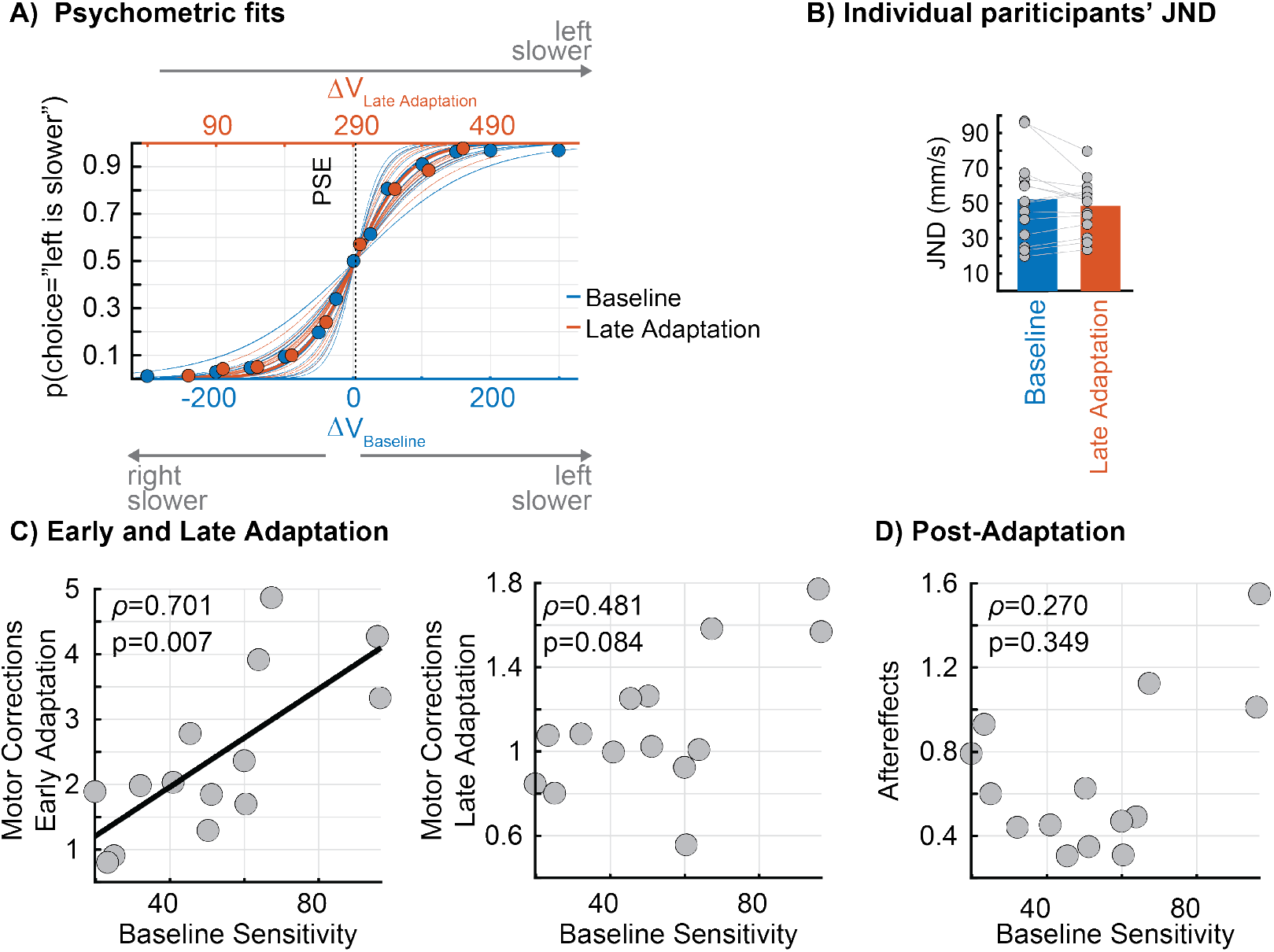
Perceptual sensitivity and its relationship to motor outcomes. **A)** Psychometric functions fitted to individual participants’ 2AFC responses during Baseline (blue) and Late Adaptation (orange), re-plotted aligned to their respective PSE values to enable direct comparison of slopes across epochs. Thin lines represent individual participants’ fitted psychometric functions and thick lines represent the group average. Circles represent pooled proportions of ‘left is slower’ responses across participants and error bars represent *±* 1.96 SE. **B)** Individual participants’ JND values at Baseline (blue) and Late Adaptation (orange). Bar height represents the group mean, gray circles represent individual participants, gray lines connect each participant’s values across epochs, and error bar represents *±* 1.96 SE. **C)** Spearman correlations between baseline sensitivity (baseline JND) and motor corrections (EMG_Magnitude_) during Early Adaptation (left) and Late Adaptation (right). Gray circles represent individual participants. Regression line is shown to indicate significant correlation. **D)** Spearman correlation between baseline sensitivity and motor aftereffects during Post-adaptation. Gray circles represent individual participants.

### Perceptual sensitivity predicts motor corrections during split-belt adaptation but not motor aftereffects

Having established that perceptual sensitivity remained the same during Baseline and Late Adaptation, we next asked whether individual differences in baseline perceptual sensitivity predicted motor corrections during Adaptation and aftereffects during Post-adaptation. Baseline JND (JND*_Baseline_*) was used as a pre-perturbation measure of individual sensory uncertainty (see Methods).

We found that baseline perceptual sensitivity was significantly associated with motor corrections during Early Adaptation (*ρ* = 0.701*, p* = 0.007; Fig 3C, left panel) but this relation was not significant during Late Adaptation (*ρ* = 0.481*, p* = 0.084; Fig 3C, right panel). Specifically, participants with greater baseline sensitivity (lower JNDs) exhibited smaller motor corrections during Early Adaptation, suggesting that perceptual sensitivity is indicative of more efficient step-by-step neuromuscular adaptation. Baseline perceptual sensitivity was not significantly associated with motor aftereffects (Fig 3D; *ρ* = 0.270*, p* = 0.349), suggesting that the benefit of high perceptual sensitivity is specific to initial motor corrections rather than the formation of motor memories. The same pattern was observed when motor corrections were quantified with step length asymmetry (perceptual sensitivity vs. step length asymmetry at early adaptation: *ρ* = 0.604*, p* = 0.025; late adaptation: *ρ* = 0.398*, p* = 0.16; aftereffects: *ρ* = *−*0.174*, p* = 0.553).

## 5 Discussion

We investigated whether perception of leg speed adapts during split-belt walking, and whether two functionally distinct dimensions of perception, perceptual prediction errors and perceptual sensitivity, relate to motor corrections and motor aftereffects. The PSE shifted toward the imposed belt-speed difference, so perceptual prediction errors were reduced, whereas perceptual sensitivity did not change between Baseline and Late Adaptation. Our findings revealed that each aspect of perception accounts for a distinct aspect of sensorimotor adaptation. Perceptual prediction errors were positively associated with motor aftereffects, whereas baseline perceptual sensitivity was associated with the initial motor corrections during Early Adaptation. Together, these findings demonstrate that perception splits into two dimensions that map onto two separate components of sensorimotor adaptation: motor memory formation on one side, and motor corrections during Early Adaptation on the other.

### Perceptual prediction error relates with motor memory formation, consistent with error-based learning

Our findings suggest that perception provides the teaching signal driving error-based learning in walking. This is novel because error-based learning has been characterized almost entirely in reduced, single-effector paradigms, where the experimenter imposes a perturbation that is assumed to directly control the prediction error driving learning (e.g., Shadmehr and Mussa-Ivaldi, 1994; Tseng et al., 2007; Lee et al., 2018). Walking affords no such control, since the experimenter sets the belt-speed difference but does not observe what the participant expects to feel, so the prediction error cannot be read off the imposed perturbation. In visuomotor adaptation, too, the error that drives implicit learning has been shown to be defined on the perceived hand direction rather than on the visible cursor that the experimenter controls (Zhang et al., 2026). We therefore used the gap between the actual belt-speed difference and each participant’s point of subjective equality (PSE), the speed difference perceived as symmetric walking, as a behavioral proxy for prediction error. If this proxy indexes the mismatch that drives learning, participants who experience larger perceptual prediction errors during adaptation should show larger aftereffects. That is what we found. Because all participants started from the same perturbation and the PSE had plateaued by Late Adaptation, we take residual perceptual prediction errors at that epoch as an index of the errors experienced during adaptation. These residual errors were positively associated with subsequent motor aftereffects, consistent with reports in reaching that larger perceptual errors accompany greater implicit adaptation (Tsay et al., 2022; Zhang et al., 2026), and with work in which error history shapes how strongly the learning system updates (Herzfeld et al., 2014; Albert et al., 2021) and in which larger or more sustained errors produce stronger motor memories (Patton et al., 2006; Scheidt et al., 2001; Thoroughman and Shadmehr, 2000; Wei and Körding, 2009; Fine and Thoroughman, 2007). Our results thus provide evidence that error-based learning extends beyond the reduced paradigms in which it was formulated to the updating of internal representations for whole-body, multi-sensory movement.

Importantly, perceptual prediction errors were quantified as a function of PSE, which represents an internal reference for what feels “normal”. This reference shifted during split-belt walking, and because a larger shift from baseline leaves a smaller gap between what feels symmetric and the actual belt-speed difference, participants with the largest perceptual shifts retained the least perceptual prediction errors at Late Adaptation. These participants showed the smallest motor aftereffects: for them, the imposed perturbation had come to feel closer to “normal”, leaving little mismatch to drive motor updating. It is therefore not the physical size of a novel perturbation alone that drives sensorimotor adaptation, but how that perturbation is perceived. In other words, the motor system may update the internal representations underlying action only to the extent that the perturbation is perceived as unusual.

The relationship we find between PSE shifts and aftereffects runs counter to studies reporting a positive association in reaching (Cressman and Henriques, 2009, 2010; Salomonczyk et al., 2011, 2013) and in an earlier walking study (Rossi et al., 2026). We believe this discrepancy reflects two distinct factors rather than a true inconsistency in the underlying phenomenon.

The first concerns how the perceptual shift relates to the perceptual prediction error in each paradigm. In visuomotor adaptation, the perturbation creates a conflict between vision and proprioception, and resolving it shifts the felt position of the hand toward the visual feedback. Participants with larger shifts in felt hand position show larger aftereffects (Cressman and Henriques, 2010; Salomonczyk et al., 2013; Ruttle et al., 2021; Tsay et al., 2021; Zhang et al., 2026), and on proprioceptive re-alignment accounts, this shift itself constitutes the error that drives implicit adaptation (Tsay et al., 2022). The split-belt perturbation, by contrast, imposes an unfamiliar mechanical condition without introducing a discrepancy between sensory modalities. Here the perceptual shift does not constitute the error but reduces it: as the PSE moves toward the imposed belt-speed difference, part of that difference comes to feel symmetric, and only the remainder is perceived as a perturbation. If a 300 mm/s difference comes to feel symmetric, for example, the imposed 500 mm/s difference is perceived as an asymmetry of only 200 mm/s. Larger PSE shifts therefore leave smaller perceptual prediction errors to drive motor adaptation, and thus smaller aftereffects. On this account, the two paradigms agree: in both, larger perceptual prediction errors accompany larger aftereffects, and the sign of the shift–aftereffect relationship differs only because the perceptual shift constitutes the error in one case and reduces it in the other.

The second factor concerns when perception and motor output are measured. We measured perceptual prediction errors at steady state, once the PSE had plateaued during Late Adaptation, which allowed us to average over trial-to-trial response variability and obtain a stable estimate within that epoch (Green and Swets, 1966; Ulrich and Miller, 2004; Ehrenstein and Ehrenstein, 1999). These errors were then related to aftereffects measured over the initial steps of post-adaptation. The earlier walking study instead assessed perceptual shifts during post-adaptation, after the perturbation had been removed (Rossi et al., 2026). In this transient phase both perception and motor output are de-adapting, so a relationship measured between them may partly reflect their shared decay, with smaller residual perceptual shifts accompanying smaller residual aftereffects, rather than a true coupling. The two designs therefore ask related but distinct questions, and relationships of the same sign need not be expected.

Our perceptual probes and Bayesian inference approach (HMM) also enabled us to estimate the evolution of perceptual shifts at the group level over the course of adaptation. The PSE shifted on average by 58% of the imposed perturbation, consistent with the partial shifts reported previously (Vazquez et al., 2015; Statton et al., 2018; Leech et al., 2018; Rossi et al., 2024, 2026). Recalibration was therefore partial. Whether this reflects a general pattern across perturbation sizes or a ceiling set by biomechanical or neural constraints remains an open question. Future studies could extend this approach to characterize subject-specific rates of perceptual adaptation and de-adaptation. Such estimates, incorporated alongside motor signals into computational frameworks of adaptation (Smith et al., 2006), could offer insight into how perception and action co-adapt over the course of learning.

### Perceptual sensitivity relates to motor corrections during Early Adaptation but not to internal model updating

Our findings suggest that perceptual sensitivity relates to motor corrections during Early Adaptation, but not to the updating of the internal models underlying movement. Bayesian integration posits that greater sensory uncertainty attenuates both feedback-driven motor corrections and feedforward recalibration, because uncertain sensory evidence is down-weighted relative to prior expectations (Körding and Wolpert, 2004; Ernst and Banks, 2002). Individuals with sharper perceptual sensitivity should therefore weight incoming sensory evidence more heavily, producing both more effective step-by-step corrections and greater recalibration of the internal models supporting feedforward control (Körding and Wolpert, 2004; Shadmehr et al., 2010; Wei and Körding, 2009; Burge et al., 2008). We found evidence for the first prediction but not the second. All participants experienced the same perturbation magnitude and maintained stability throughout, yet those with greater baseline sensitivity did so with smaller muscle activity during early adaptation, indicating that they counteracted the perturbation more efficiently, since reduced muscle activity during walking corresponds to lower metabolic cost (Finley et al., 2013; Sánchez et al., 2019). This is consistent with work showing that higher sensitivity to external perturbations is associated with better motor responses for maintaining balance in young adults (Mirdamadi et al., 2024) and with faster adaptation in older adults (Lei and Wang, 2018).

In contrast, baseline sensitivity was not associated with motor output during late adaptation and post-adaptation. Dual-rate models of motor adaptation offer an explanation (Smith et al., 2006): a fast, reactive process dominates early on, while a slow, feedforward process gradually takes over and eventually produces aftereffects. As the feedforward process takes over, motor output increasingly reflects stored internal-model updates, and feedback-mediated corrections matter less. On this account, our results suggest that perceptual sensitivity bears on feedback control rather than on the recalibration of internal models, in line with null findings from reaching tasks (Vandevoorde and Orban de Xivry, 2021; Kitchen and Miall, 2021), which we now extend to continuous, multi-sensory locomotor behavior. In visuomotor rotation, lower proprioceptive variability, that is, greater sensitivity, has been reported to predict less implicit adaptation (Tsay et al., 2021, 2022), which is consistent with Bayesian integration because the perturbation there is visual, and greater proprioceptive sensitivity reduces the weight given to the perturbed visual signal (Tsay et al., 2021). Replication in a larger sample is needed before the absence of a relationship is taken as evidence against a role for Bayesian weighting in feedforward recalibration.

### Split-belt walking recalibrates perceptual bias without changing sensitivity

Perceptual sensitivity remained the same between Baseline and Late Adaptation even as the PSE, the internal reference of what feels symmetric, shifted, indicating that the mechanisms recalibrating bias and those determining sensitivity are at least partially independent. The same pattern appears in visuomotor adaptation (Cressman and Henriques, 2009; Salomonczyk et al., 2013) and force-field paradigms (Kitchen and Miall, 2021; Ostry et al., 2010), suggesting a general organizational principle of sensorimotor recalibration rather than a feature specific to upper-limb or visually guided tasks.

Framed in Bayesian terms (Körding and Wolpert, 2004), this dissociation suggests that adaptation selectively updates the belief over body and environmental states without altering the precision of the likelihood. The JND, as a behavioral index of that precision, reflects internal sensory measurement noise, which Bayesian observer models separate from the prior expectations that set perceptual biases (Stocker and Simoncelli, 2006). The dissociation between a shifting PSE and an unchanged sensitivity was only detectable because we used a 2AFC paradigm, whereas most prior studies used speed-matching (Vazquez et al., 2015; Statton et al., 2018; Leech et al., 2018; Rossi et al., 2026) or single-threshold tasks (Lauzière et al., 2014; Hoogkamer et al., 2015), which yield a single perceptual estimate akin to the PSE and cannot assess whether sensitivity also changes. In 2AFC paradigms, reaction times are also available, and they could be used to estimate perceptual sensitivity separately from the choices (Gonzalez-Rubio et al., 2025). Reaction times contain information about stimulus strength beyond that contained in choices, and the two can be modeled jointly (Stone, 2014), which may allow sensitivity to be estimated from fewer trials and shorten the perceptual probes during adaptation.

### EMG_Magnitude_ and step length asymmetry differ in their sensitivity to perception-action relationships

We chose EMG activity as our primary motor outcome because it has been shown to be more sensitive to individual differences in sensorimotor recalibration than kinematic measures such as step length asymmetry (Iturralde and Torres-Oviedo, 2019). When step length asymmetry was used as a secondary motor outcome measure, the association between baseline sensitivity and motor corrections during Early Adaptation was recovered (*ρ* = 0.604, *p* = 0.025), but the association between perceptual prediction errors and motor aftereffects was not. One possibility is that this reflects the temporal resolution of the two measures: EMG_Magnitude_ is computed across the entire gait cycle, whereas step length asymmetry summarizes a single time point within it, so individual differences in perception-action coupling distributed across the cycle would not be captured by a discrete kinematic metric. Consistent with this, movement quality quantified from full kinematic trajectories has been shown to be more sensitive than features sampled at single time points for detecting differences in motor control across individuals (Arac et al., 2026). Furthermore, EMG_Magnitude_ aggregates the activity of multiple muscles, so it may better capture individual differences in adaptation. Namely, the same step lengths can be produced by different patterns of muscle activity, so individual differences in adaptation may appear in muscle activity without appearing in step length asymmetry. Future work using continuous kinematic trajectories, and muscle-recruitment patterns in high-dimensional muscle space, could test these possibilities directly and clarify how perceptual dimensions relate to the coordination (Iturralde and Torres-Oviedo, 2019), rather than only the magnitude, of motor patterns.

### Implications

Our findings generate specific, testable predictions for populations in which motor adaptation is differentially affected. Individuals post-stroke show increased motor corrections and slower adaptation (Tyrell et al., 2015; Savin et al., 2013) yet retain the ability to form aftereffects comparable to controls (Tyrell et al., 2015; Savin et al., 2013), despite common lower-limb proprioceptive deficits (34–64% of survivors; Connell et al. 2008; Dukelow et al. 2010; Chia et al. 2019; Tyson et al. 2008). Our framework predicts that these motor deficits may reflect impaired perceptual sensitivity specifically, while intact aftereffects would be consistent with preserved perceptual prediction error minimization and internal model updating.

In healthy aging, older adults show steady-state kinematic adaptation comparable to young adults (Aucie et al., 2021; Roemmich et al., 2014), but findings on aftereffects are mixed, reduced in some studies (Iturralde and Torres-Oviedo, 2019; Bruijn et al., 2012) and comparable or larger in others (Sombric et al., 2017; Swart et al., 2023), despite well-documented declines in lower-limb proprioception with age (Petrella et al., 1997; Thelen et al., 1998; Yang et al., 2019). Our framework predicts that if older adults retain sufficient sensitivity to detect inter-leg speed differences, their feedback-driven corrections should remain intact, consistent with generally preserved adaptation performance, whereas declines in aftereffects would instead point to feedforward impairments; the variability across studies may reflect heterogeneity in how each pathway is affected by aging, physical activity level (Hiew et al., 2023), or task demands.

Individuals with cerebellar damage are especially informative, given the cerebellum’s role in feedforward control and internal model updating (Wolpert et al., 1998; Bastian, 2006; Tseng et al., 2007). During split-belt walking, cerebellar patients show intact early corrections but fail to reduce asymmetry over adaptation and produce little to no aftereffects (Morton and Bastian, 2006), a deficit that scales with ataxia severity (Statton et al., 2018), despite intact ability to detect belt-speed differences (Morton and Bastian, 2006), that is, intact sensitivity alongside absent error-driven learning. Cerebellar damage also selectively impairs active but not passive proprioception, consistent with disrupted predictive modeling rather than degraded peripheral sensation (Bhanpuri et al., 2013). This pattern aligns with the dissociation we observed in healthy young adults, where sensitivity supports corrections that do not depend on cerebellar integrity, while perceptual prediction error tracks the feedforward process that does. Directly measuring both dimensions in cerebellar, stroke, and aging populations with the paradigm used here would test whether this dissociation generalizes, and could help localize the source of adaptation deficits across clinical populations.

## 6 Methods

### 6.1 Data Collection

#### Participants

Fourteen neurotypical young adults (9 females, 20.86 *±* 1.92 y.o.) participated in this study. Two of the participants reported left-handedness, and three reported being left-footed when kicking a ball (Kramer and Balsor, 1990). All participants were naïve to split-belt walking. The protocol was approved by the University of Pittsburgh’s Internal Review Board (IRB) in accordance with the Declaration of Helsinki, and all participants gave written informed consent prior to participation.

#### Perceptual task

We assessed participants’ ability to detect interlimb leg-speed differences through a 2-alternative forced choice task (refer to Fig 1A), like the one implemented in Gonzalez-Rubio et al. (2025). Initially, subjects walked on a set speed difference between the legs that was either set to zero or 500 mm/s depending on the protocol epoch where we assessed perception. Then, participants experienced a sudden introduction of a stimulus, which is defined as a difference in speed between the right (R) and left (L) leg (Δ*V* = *V_R_ − V_L_*). This asymmetry was created by accelerating one treadmill belt while decelerating the other in sequence beginning with the right side, always maintaining the average speed between the two legs. Speed changes on each belt occurred during the swing phase when the corresponding foot was in the air. An auditory signal marked the onset of the decision period (indicated by gray shading in Fig 1), which was limited to a maximum of 8 seconds (approximately 8 strides). Upon this start cue, participants used handheld devices to indicate their perception of which limb was slower by pressing a key on either the right or left controller. No indication was given to prioritize either response speed or accuracy. If no response was recorded, a second, distinct auditory warning cue was delivered to indicate 2 seconds left in the task and prompt participants to register a response. The perceptual task concluded upon response registration or if no response was given within the allotted time. Following their response, participants received an auditory message indicating their selection (e.g.,“Left is slow”), ensuring they were aware their response had been recorded. Notably, this message did not provide any information on the accuracy of their response.

All auditory cues were transmitted via noise-canceling headphones, including start, response, and warning signals. To isolate somatosensory processing from other sensory inputs, we implemented white noise, masking auditory signals, and visual occlusion of the lower extremities using an opaque barrier, eliminating potential environmental cues that might bias perceptual judgments.

#### Experimental design

The overall protocol is outlined in Figure 1B. Each participant completed two sessions of data collection on separate days (i.e., within a week). Each experimental session started with a Familiarization period consisting of 4 perceptual tasks practice trials. During this phase, participants received real-time visual feedback through dynamic belt speed displays on a TV monitor as well as guidance from research staff. This stage guaranteed participant comprehension and comfort with experimental procedures prior to data collection. For the first session, following familiarization, participants completed 3 experimental blocks of data collection (Fig 1B, top panel). Within each block, participants completed 56 perceptual tasks encompassing each non-zero stimulus magnitude (Δ*V* could take any value among *±*25, *±*50, *±*100, *±*150, *±*200, and *±*300 mm/s) where each was presented 4 times within a block, alongside null perceptual tasks (0 mm/s) administered 8 times. Thus, each non-zero stimulus magnitude was repeated 12 times, and the zero stimulus (null perceptual task) was repeated 24 times. Importantly, between perceptual tasks, participants walked with both belts moving at the same speed (1.05 m/s). Stimuli were presented in pseudorandom sequence, with identical ordering maintained across all subjects. We stopped the treadmill between blocks, and once within, providing 3-minute rest periods to prevent cognitive and physical fatigue.

In Session 2 (Fig 1B, bottom panel), following the Familiarization period, participants completed a brief baseline assessment similar to what they experienced in Session 1; however, they only had two presentations of the *±*100, *±*200, and *±*300 mm/s stimulus magnitudes, and four presentations of null perceptual tasks (0 mm/s) in a single block. We selected this subset of non-zero stimulus magnitudes because these were the ones that participants experienced initially in the adaptation period (as described below). During the Adaptation period, all perceptual tasks were interleaved every 16 strides of a sustained belt-speed difference of 500 mm/s, with each participant’s dominant leg positioned on the faster belt. Perceptual trials followed two distinct schedules between the “Transient” and “Late Adaptation” phases of adaptation to track more accurately the changes in perception. During the “Transient” phase, perceptual tasks began after 10 strides of split-belt walking. The stimuli magnitudes progressed through speed differences of +100 and +200 mm/s, then +200 and +300 mm/s, and finally +200 and +400 mm/s. Within each pair of stimulus magnitudes, the two stimulus were presented in alternating order 5 times per pair (i.e., 10 trials per pair, 30 trials total). During the “Late Adaptation” phase perceptual tasks spanned stimuli magnitudes ranging from +50 to +450 (in 50 mm/s increments) with 10 repetitions of each stimuli magnitude presented in pseudo-random order. We selected these stimuli magnitudes based on pilot data indicating the point of subjective equality would on average shift to approximately 250 mm/s by late adaptation. This “Late Adaptation” phase of adaptation had more repetitions of each stimuli magnitude to fit logistic functions as done in our prior studies (Gonzalez-Rubio et al., 2025). The “Transient” and “Late Adaptation” phases of adaptation encompassed a total of 120 perceptual tasks interspersed with approximately 2150 strides of split-belt walking over 60 minutes. Five scheduled 3-minute breaks were provided, with additional rest periods available upon request. During the Post-adaptation period, perceptual tasks began after 10 strides of tied-belt walking. The stimulus magnitudes progressed through +200 and +150 mm/s, then +50 and +100 mm/s, where each of the two stimulus pairs were presented twice in an alternating order. Finally, the remaining trials varied between 0, *±*50, and *±*100 mm/s in pseudo-random, for a total of 22 perceptual tasks.

The first perceptual task after each resting break was excluded from the analysis, and an additional trial of the same magnitude was appended before the next resting break. This ensured that participants were not caught off guard, which could have led to unintended non-responses. Also, non-response tasks were excluded from the analysis (a total of 17 out of 4562 perceptual tasks), corresponding to 0.37% of all perceptual tasks.

### 6.2 Data Analysis

#### Perceptual task outcomes

From each 2AFC task, we extracted participants’ choice responses (indicating which leg was perceived as slower). This measure was recorded for all tasks, including null perceptual tasks of 0 mm/s stimulus magnitude.

#### Point of subjective equality and just noticeable difference

Using the choice data, we calculated two key perceptual measures: the point of subjective equality (PSE) and just noticeable difference (JND). The PSE represents the stimulus magnitude where the probability to choose either ‘left’ or ‘right’ is 50%, this metric indicates the interlimb leg-speed that participants perceive as being symmetric. We were interested in quantifying the perceived interlimb speed symmetry because it has been used to assess perceptual aftereffects following split-belt walking (Vazquez et al., 2015; Statton et al., 2018; Leech et al., 2018; Rossi et al., 2026, 2024) and it has been related to motor aftereffects (Rossi et al., 2026)

The JND quantifies the minimum stimulus difference reliably detected by participants. Following established psychophysical methods (Bush, 1963; Paire et al., 2023; Aida et al., 2020; Bausenhart et al., 2012; Ulrich, 2010), we defined JND as half the distance between the stimuli producing 25% and 75% choice rates. This approach captures participant sensitivity both below and above the PSE by averaging two sensitivity measurements: the difference between 25%-50% choice rates and between 50%-75% choice rates, providing a comprehensive assessment of perceptual sensitivity across the stimulus range (Ulrich and Vorberg, 2009). Since empirical data rarely yield exact 25%, 50%, and 75% choice rates, we used psychometric curve fitting to interpolate these values and calculate PSE and JND accordingly.

#### Psychometric curve fitting

We characterized the relationship between participants’ choices (left or right) and the stimulus magnitude (Δ*V*) for the 2AFC tasks, following established methods from Gonzalez-Rubio et al. (2025). The probability of a participant choosing “left” was represented using a logistic function of *µ_j_*, where *j* denotes individual participants, as described in Eq. 1.

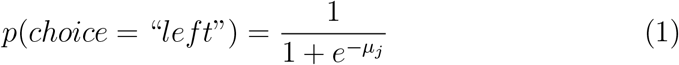

#### Testing adaptation of perception

To determine whether perception of speed differences after split-belt walking changes, we formulated our generalized linear mixed model with *µ_j_* defined as follows:

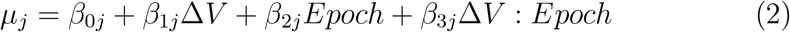

Where *Epoch* refers to the experimental phase (baseline or late adaptation) where we had enough perceptual tasks to generate full psychometric functions. This categorical variable takes the value of zero when referring to perception during baseline (evaluated during Session 1) and 1 when referring to perception during late adaptation (Session 2, with the “Late Adaptation” perceptual tasks). Here, *β*_1_*_j_* and *β*_1_*_j_* + *β*_3_*_j_* captures the sensitivity to stimulus (Δ*V*) for the baseline and late adaptation epochs respectively. Similarly, the bias can be thought of *β*_0_*_j_* and *β*_0_*_j_* + *β*_2_*_j_*. Accordingly, *β*_2_*_j_* and *β*_3_*_j_* indicates a change between the two epochs.

All models included participant as a random effect to account for individual differences in both intercept and slope parameters. The fixed effect estimates (*β̄*_0_, *β̄*_1_, *β̄*_2_, and *β̄*_3_) represent average parameters across participants. Model fitting was performed using MATLAB’s (The Mathworks, Inc., Natick, Massachusetts, United States) *fitglme* function with maximum likelihood estimation using the Laplace approximation method. Statistical significance of the epoch effects (*β̄*_2_ and *β̄*_3_) was assessed using Wald tests (*H*_0_ : *β* = 0).

PSE and JND were directly estimated from the regressors from Eq. 1 and 2. Specifically, the relationship between the model parameters and the PSE for each participant *j* and each epoch is:

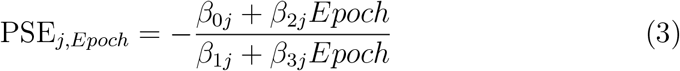

The relationship between the model parameters and the JND for each participant *j* and each epoch is:

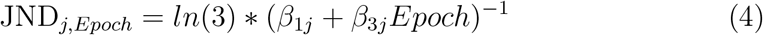

We also performed paired t-tests between the Baseline and Late Adaptation PSE and JND values to directly compare these derived measures across epochs. The significance level was set to *α* = 0.05.

#### Perceptual outcome measures for regression analyses

We related perceptual adaptation to motor adaptation using two perceptual outcome measures: perceptual prediction error and perceptual sensitivity. For each participant *j*, perceptual prediction error was defined as the difference between the imposed belt-speed difference (500 mm/s) and the shift in the point of subjective equality (PSE) from Baseline to Late Adaptation:

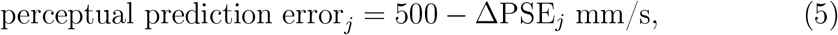

where ΔPSE*_j_* = PSE*_j,_*_LateAdaptation_ *−* PSE*_j,_*_Baseline_. Thus, a larger perceptual prediction error indicates that a participant’s sense of symmetry shifted less toward the imposed perturbation. Perceptual sensitivity was quantified as each participant’s Baseline JND (JND*_j,_*_Baseline_), where smaller JND values indicate greater sensitivity.

#### Acquisition and pre-processing of kinematics, kinetic, and muscle activity data

We collected kinematics, kinetic, and electromyography (EMG) signals data to characterize participants’ motor behavior. Kinematic data were collected at 100 Hz using a 14-camera motion analysis system (Vicon Motion Systems). We recorded movement by placing markers bilaterally on anatomical landmarks including the lateral malleolus (ankle) and greater trochanter (hip). We also recorded ground reaction forces from our instrumented splitbelt treadmill (Bertec Corporation) and sampled the data at 1000 Hz. We used the ground reaction forces along the vertical axis to determine gait cycle events using a threshold of 30 N. Step length was computed as the anteriorposterior distance between the ankle markers at heel strike (Finley et al., 2015), and we computed step length asymmetry as the difference between fast and slow leg step lengths, normalized to their sum. Surface EMG electrodes were placed on 5 muscles of the lower legs: tibialis anterior, peroneus longus, soleus, and the lateral and medial gastrocnemius. These EMG signals were sampled at 2000 Hz using the Delsys Trigno System (Delsys Inc.). EMG signals were high-pass filtered to remove movement artifacts, rectified, and then filtered using a second-order Butterworth filter with dual-pass filtering and a cutoff frequency of 30 Hz. The filtered EMG activity was aligned to gait events and divided into 12 intervals for higher temporal resolution (Iturralde and Torres-Oviedo, 2019). Each muscle’s activity was first normalized to its maximum baseline activity, making it dimensionless and expressed as a percentage of baseline. The magnitude of muscle activity for each stride t (EMG_t_) was then computed as the Euclidean norm of the resulting 120dimensional vector (10 muscles x 12 sub phases of the gait cycle; Iturralde and Torres-Oviedo 2019).

#### Motor outcome measures for regression analyses

The magnitude of muscle activity for each subject was used to compute three outcome measures: motor corrections during early and late adaptation and motor aftereffects. These three outcome measures were used to assess the relationship between perceptual and motor adaptation. We computed motor corrections during Early Adaptation as the mean magnitude of muscle activity EMG_EarlyAdaptation_ across the first 30 strides of the Adaptation epoch. Motor corrections during Late Adaptation were computed as the mean magnitude of muscle activity EMG_LateAdaptation_ across the last 40 strides of split-belt walking during Adaptation epoch (excluding the very last 5 strides) to quantify the “efficiency” of neuromuscular motor patterns at the end of adaptation. In both epochs, smaller values indicate less muscle activity, which we refer to as more efficient motor corrections. Lastly, motor aftereffects were computed as the averaged magnitude of muscle activity EMG_Aftereffects_ across the first 10 strides of post-adaptation to quantify the strength of motor recalibration when the split-belt perturbation is removed.

#### Secondary motor outcome measures

As a secondary analysis, we computed analogous motor outcome measures using step length asymmetry, the conventional kinematic measure used during split-belt adaptation (Reisman et al., 2007; Morton and Bastian, 2006; Torres-Oviedo and Bastian, 2012; Finley et al., 2015). Step length asymmetry during early adaptation was computed as the mean across the first 30 strides of the Adaptation epoch, late adaptation as the mean across the last 40 strides (excluding the final 5 strides), and aftereffects was the mean across the first 10 strides of Post-adaptation, matching the stride windows used for the EMG_Magnitude_. We selected EMG_Magnitude_ as our primary motor outcome measure because it has been shown to be more sensitive to individual differences in sensorimotor recalibration than step length asymmetry (Iturralde and Torres-Oviedo, 2019).

#### Relationship between perception and action

To investigate the relationship between the perceptual and motor domains, we used Spearman rank correlations (*ρ*) to examine whether the magnitude of perceptual prediction error was associated with motor correction measures (EMG_Magnitude_ during Early Adaptation, Late Adaptation, and Aftereffects). We then used a second set of correlations to determine if baseline perceptual sensitivity (JND_Baseline_) was associated with motor corrections. Spearman correlations were chosen because they do not assume linearity or normality and are robust to outliers, which is appropriate given our sample size. Separate correlations were computed for each motor outcome measure at each epoch. The significance level was set to *α* = 0.05, corrected for multiple comparisons across the three epochs using the Bonferroni method (*α*_corrected_ = 0.0167).

### 6.3 Markov Modeling

To estimate the continuous evolution of the PSE throughout the protocol, we employed a Hidden Markov Model (HMM). The latent state *x_t_* represents the PSE (in mm/s) at stride *t*, evolving as a Gaussian random walk (*x_t_*= *x_t−_*_1_ + *w_t_*, *w_t_∼ N* (0*, σ*^2^)). Binary 2AFC responses were modeled using a logistic psychometric function with lapse rate:

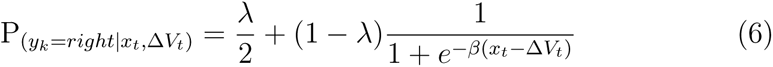

where Δ*V_t_* is the belt-speed difference at stride *t*, *β* is the perceptual slope, and *λ* is the lapse rate. This follows the same logistic psychometric model used to characterize perceptual sensitivity in our main analysis, but differs in two aspects: first, here we fit a standard maximum likelihood estimate on the pooled data without considering random mixed effects; second, the model includes a lapse rate parameter *λ* to account for attentional lapses or random responses (Treutwein and Strasburger, 1999; Wichmann and Hill, 2001). The psychometric parameters *β* and *λ* were estimated from the data collected in Session 1 (Baseline) data by fitting a logistic psychometric function to the pooled 2AFC responses across all participants via maximum likelihood, yielding *β* = 0.027 and *λ* = 0.047 (JND = 40.8 mm/s). These values were held fixed for the HMM applied to Session 2.

Because the Bernoulli observation model precludes closed-form posterior inference, the continuous PSE was discretized on a fine uniform grid (Kitagawa, 1987), and smoothed posterior distributions were obtained via the forward–backward algorithm (Baum et al., 1970; Rabiner, 1989). The prior was initialized at PSE = 0 mm/s. The transition noise *σ* = 4.5 was selected from a grid of candidate values by maximizing the marginal log-likelihood of the observed data. Observations were aggregated across participants to estimate a single group-level PSE trajectory. Each block was fitted as an independent HMM pass, that is, the smoothed posterior at the end of one block was propagated forward through the inter-block gap using the transition model, allowing uncertainty to grow during unobserved walking, and the resulting distribution served as the prior for the next block. This approach avoids imposing an arbitrary starting point for each block and instead lets the observations determine the PSE estimate. Apparent discontinuities at block transitions reflect the increased uncertainty accumulated during the gap, the broad prior is rapidly updated by incoming observations, which may shift the estimate.

## Data and code availability

The dataset and code used during the current study are available on Open Science Framework (https://osf.io/3xt5b/).

## Acknowledgments

This research was supported in part by the U.S. National Science Foundation (CAREER Award 1847891 to G.T.O. and M.G.R., Award 2419849 to G.T.O. and M.G.R., and Graduate Research Fellowship Program Award 2139321 to M.G.R.) and by the National Institutes of Health (Award R90DA060340-02S1 to G.T.O. and M.G.R.). The funders had no role in study design, data collection and analysis, decision to publish, or preparation of the manuscript.

## Author contributions

M.G.R., G.T.O., and P.A.I. were involved in the conception and design of the research. M.G.-R. and A.R.C. performed the research. M.G.R., A.R.C., G.T.O., and P.A.I. analyzed the data and interpreted the results. M.G.R., G.T.O., and P.A.I. wrote the manuscript. The final version of the manuscript was approved by all authors, who agree to be accountable for all aspects of the work in ensuring that questions related to the accuracy or integrity of any part of the work are appropriately investigated and resolved. All authors qualify for authorship, and all those who qualify for authorship are listed.

